# Spatiotemporal Dynamics of Highly Pathogenic Avian Influenza H5 Virus Introductions and Regional Spread in the Republic of Korea

**DOI:** 10.64898/2026.05.21.726857

**Authors:** Taehee Chang, Sangyi Lee, Jin Il Kim, Kyung-Duk Min

## Abstract

Highly pathogenic avian influenza (HPAI) viruses from clade 2.3.4.4 have caused recurrent outbreaks in poultry since 2014. In the Republic of Korea, clade 2.3.4.4b viruses have driven five epidemic waves, yet the factors underlying HPAI introduction and farm-to-farm spread remain poorly understood. We compiled hemagglutinin gene sequences of clade 2.3.4.4b viruses from wild birds and poultry in the Republic of Korea (October 2016–March 2024) and reconstructed dispersal dynamics using Bayesian phylogeography. Dispersal patterns suggest that domestic duck farms in the western provinces likely form a key interface for spillover from wild birds into poultry. Mixed-effects generalized linear models showed that both wild-to-poultry and farm-to-farm transition rates were positively associated with the number of poultry farms in the destination province, while wild-to-poultry rates were further associated with higher avian influenza virus infection probability among wild birds. Wild-to-poultry transition rates were lower in 2020–2024 than in 2016–2018, which may reflect strengthened interventions. These findings suggest that poultry farm abundance and introduction pressure from wild birds jointly shape the spatial dynamics of HPAI introduction and spread. More broadly, these factors may provide operational indicators to guide risk-based surveillance and control strategies.

**Author Summary:** Highly pathogenic avian influenza (HPAI) H5 viruses continue to cause major losses in poultry and pose recurring risks at the wildlife–livestock interface. Effective control depends on identifying where viruses are most likely to enter poultry populations and how they spread between farms. Using viral genomic data from wild birds and poultry in the Republic of Korea, this study suggests that domestic duck farms in western provinces likely form a key interface for introductions from wild birds into poultry. We also found that regions with more poultry farms were more likely to receive and spread the virus, while introduction risk was further elevated where infection pressure from wild birds was higher. By linking viral genomic patterns with ecological and epidemiological information, our study helps identify where HPAI viruses are most likely to enter poultry populations and spread between farms. These findings can guide targeted surveillance and early control in regions at greatest risk.

## 1. INTRODUCTION

Understanding the transmission dynamics of zoonotic viruses within animal reservoirs is critical for preventing spillover into human populations (1). Influenza A viruses (genus Alphainfluenzavirus, family Orthomyxoviridae) represent a major zoonotic threat due to their extensive genetic diversity, broad host range in avian reservoirs, and repeated spillover events to mammals, including humans (2). Importantly, these viruses pose a significant risk of causing a human influenza pandemic (2).

Wild aquatic birds are the primary reservoir hosts for avian influenza A viruses (AIVs), where diverse viral lineages circulate mainly in weakly pathogenic form (3). In poultry, however, these viruses can evolve into highly pathogenic avian influenza (HPAI) through insertions at the hemagglutinin (HA) cleavage site, enabling systemic infection (4). One example is the goose/Guangdong (Gs/Gd) lineage HPAI H5N1 virus, which emerged in China in 1996 and has since become endemic in domestic poultry (5). Following its emergence, Gs/Gd viruses diversified through mutations in HA, giving rise to multiple antigenically distinct clades (6). Among these, clade 2.3.4.4 (a–h) has undergone extensive reassortment, generating H5Nx variants with neuraminidase (NA) subtypes other than N1 (7).

Since 2014, spillback events of clade 2.3.4.4 viruses from poultry to wild birds have facilitated episodic long-distance dissemination, contributing to recurrent outbreaks in poultry and wild birds across five continents (8–10). Since 2016, H5N8 viruses from the clade 2.3.4.4b have been implicated in repeated outbreaks among wild birds (11). Since November 2021, reassortant H5N1 viruses from clade 2.3.4.4b have largely replaced H5N8 from the same clade and have driven unprecedented outbreaks in poultry and wild birds. Sustained circulation in Europe and the Americas, regions previously considered epidemiological sinks, has raised concerns that endemic establishment may expand geographically (12–14). In addition, clade 2.3.4.4 viruses have increasingly been associated with spillover into both wild and domestic mammals (10, 15–17). Although the genetic origins and large-scale dissemination of Gs/Gd viruses have been extensively documented (10, 18–20), the key drivers of AIV introductions from wild birds to poultry and subsequent farm-to-farm transmission remain insufficiently understood. This knowledge gap constrains risk-based control efforts in both endemic and non-endemic regions that experience episodic incursions.

The Republic of Korea (ROK) has experienced repeated introductions of Gs/Gd viruses leading to outbreaks on poultry farms, with 11 epidemic waves recorded from the first detection in December 2003 through spring 2024 (21, 22). Of these, five waves since November 2017 were caused by clade 2.3.4.4b viruses, including clade 2.3.4.4b H5N6 during 2017–2018 (23), clade 2.3.4.4b H5N8 during 2020–2021 (24), and clade 2.3.4.4b H5N1 during 2021–2022 (25–27), 2022–2023 (21, 26, 28, 29), and 2023–2024 (22, 30). Together, these five waves accounted for a cumulative total of 264 reported farm-level outbreaks. Although these waves were generally smaller than the catastrophic 2016–2017 epidemic (31, 32), HPAI outbreaks have continued to occur on poultry farms despite the implementation of enhanced and reorganized national biosecurity systems (33, 34). This pattern highlights the urgent need to strengthen risk-based surveillance and control measures to prevent recurrent AIV introductions and subsequent farm-to-farm spread.

Although outbreaks have occurred nationwide, they have been disproportionately concentrated in the western coastal provinces of the Korean Peninsula, where there are many wintering sites for migratory birds and a high density of duck farms (23, 34). Regardless of AIV subtype or clade, early affected farms tend to cluster near locations where AIV has been detected in wild birds (35, 36). This suggests spatially structured heterogeneity in AIV incursion risk for poultry farms, likely shaped by interplay among the movements of migratory birds (23, 33), local environmental conditions (34), and anthropogenic factors such as high poultry density and farm-to-farm connectivity (13). Accordingly, mapping how viruses enter and spread between farms, while pinpointing the ecological and epidemiological factors that drive these processes, can help direct resources toward risk based interventions.

In this study, we characterized the dispersal patterns of clade 2.3.4.4b viruses associated with recurrent epidemic waves in the ROK. We compiled HA gene sequences of these viruses sampled from wild birds and poultry in the ROK between October 2016 and March 2024, spanning five epidemic waves. Across these waves, we compared dispersal patterns between early and recent epidemics. We reconstructed spatiotemporal dispersal dynamics between key host groups and across administrative regions. By leveraging Bayesian phylogeographic approaches, we resolved directional pathways of AIV transmission, distinguishing introductions from wild birds into poultry and subsequent farm-to-farm spread, thereby complementing insights from previous analyses. We also evaluated ecological and epidemiological factors associated with inferred transition rates.

## 2. RESULTS

### 2.1. Spatiotemporal patterns of HPAI H5 outbreaks

In the ROK, five epidemic waves associated with HPAI H5 viruses were observed between October 2017 and March 2024 (Figure S1). In total, 264 farm-level outbreaks were recorded (Table 1). Spatially, outbreaks were concentrated along the western coast and showed spatial heterogeneity by poultry host group (Figure 1A and S2). This heterogeneity is not fully explained by the underlying distribution of farms. To account for differences in the number of farms, we calculated standardized incidence ratios (SIRs; observed-to-expected ratios) for farm-level outbreaks. SIRs exceeded 1 for domestic Galliformes farms (primarily chickens) in Gyeonggi-do (GG) and for domestic Anseriformes farms (primarily ducks) in Jeollabuk-do (JB) and Jeollanam-do (JN) (Table 1). Temporally, outbreaks typically began in October, peaked in December, and subsided by March (Figure 1B). Monthly patterns further suggested an earlier onset in domestic Anseriformes farms in JB and JN, followed by increased activity in GG and other provinces. In addition, the HPAI H5 viruses circulating in the ROK during this period were inferred to belong predominantly to clade 2.3.4.4b, and multiple subtypes (e.g., H5N6, H5N8, and H5N1) appeared to circulate either sequentially or concurrently (Figure S3). The MCC phylogeny inferred from sequences from the ROK showed clear clustering by H5 HA clades (Figure S4).

**Figure 1.**
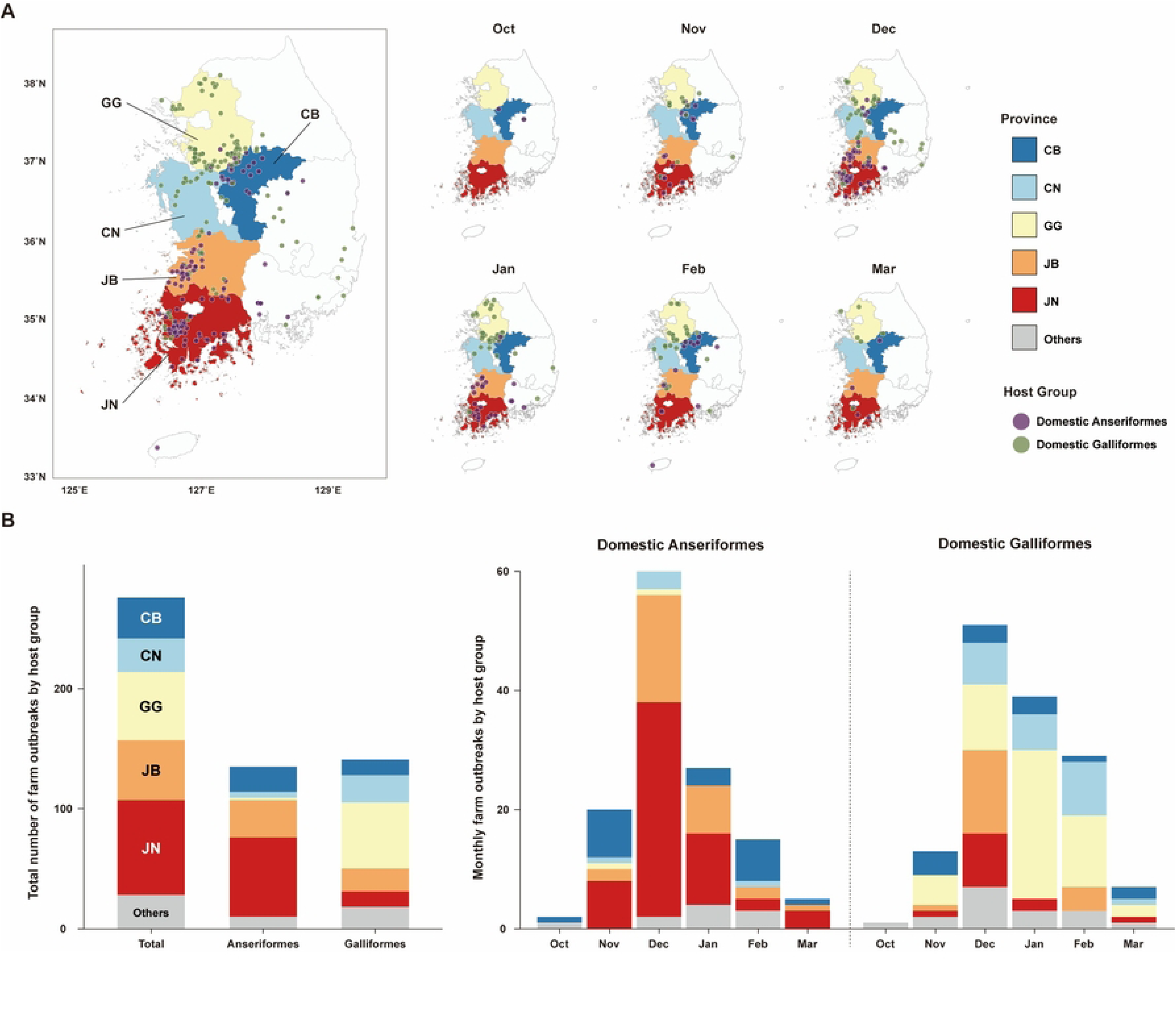
Spatiotemporal patterns of outbreaks on poultry farms. (A) Faceted maps showing the locations of affected farms by month (October–March), with counts pooled across all epidemic seasons (October 2017–March 2024). Points indicate outbreak locations and are colored by poultry type. (B) Stacked bar charts summarize the total number of reported outbreaks on poultry farms by month (October–March). Within each bar, colored segments represent province-specific outbreak counts, stacked to sum to the monthly total (*y*-axis). Counts for domestic Anseriformes (primarily ducks) and domestic Galliformes (primarily chickens) are shown in separate panels. Provinces with the five highest outbreak counts are shaded.

**Table 1.**
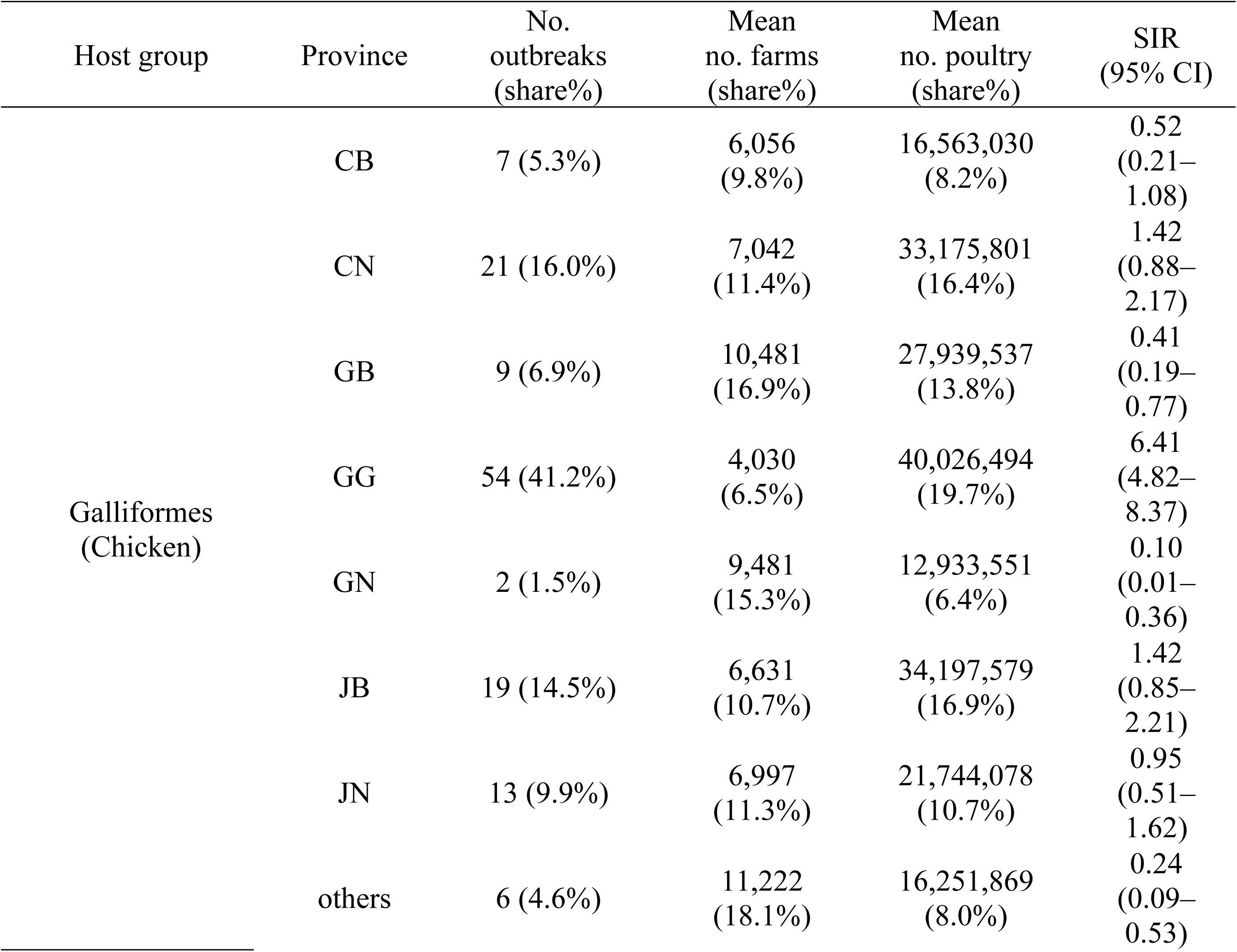
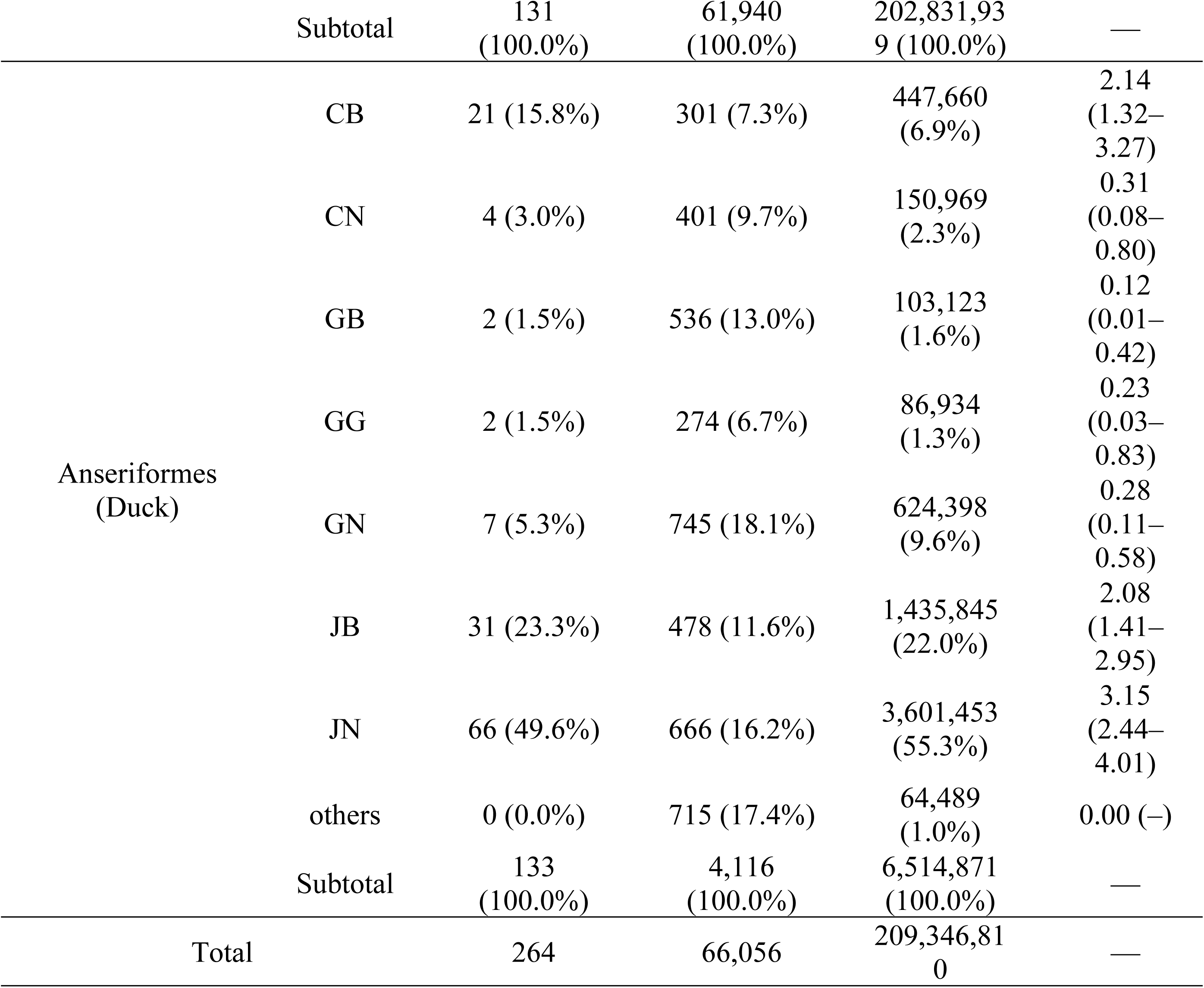
Regional distribution of outbreak events in poultry flocks and production metrics by host group. Outbreak counts represent the total number of farm-level outbreak events reported in monthly records covering October 2017 through March 2024 and restricted to outbreaks associated with HPAI viruses. Farm counts and poultry population (head count) correspond to province-level annual values summarized as the mean for 2017–2023 (annual data for 2024 were unavailable). Percent shares were calculated within each host group. SIRs denote the observed-to-expected outbreak ratio (O/E); 95% confidence intervals were calculated assuming Poisson-distributed observed counts. Provinces are grouped as CB, CN, GB, GG, GN, JB, and JN, while all remaining provinces are aggregated as “Others.” Subtotals summarize values within each host group, and the overall total represents the sum across host groups.

### 2.2. Host transition dynamics

To characterize inter-host transition dynamics of clade 2.3.4.4b in the ROK, we reconstructed host-trait transitions along the phylogeny using a discrete-trait continuous-time Markov chain model (Figure 2 and S5). Host-transition pathways with strong statistical support (Bayes factor [BF] > 10) were inferred for transitions from wild birds to domestic Anseriformes and from domestic Anseriformes to domestic Galliformes. Transitions from domestic Anseriformes to wild birds and from wild birds to domestic Galliformes received moderate support (3 < BF ≤ 10). Markov reward analysis suggested that viral lineages have been maintained for longer evolutionary periods in domestic Anseriformes than in domestic Galliformes. In addition, inferred wild-to-poultry transitions exhibited seasonality, peaking in November or December and declining through March. Overall, these patterns suggest that domestic Anseriformes likely act as an intermediary in the transmission of 2.3.4.4b viruses from wild birds to poultry farms, with such transitions occurring more frequently during the early phase of the epidemic season. These patterns were broadly consistent across analysis windows.

**Figure 2.**
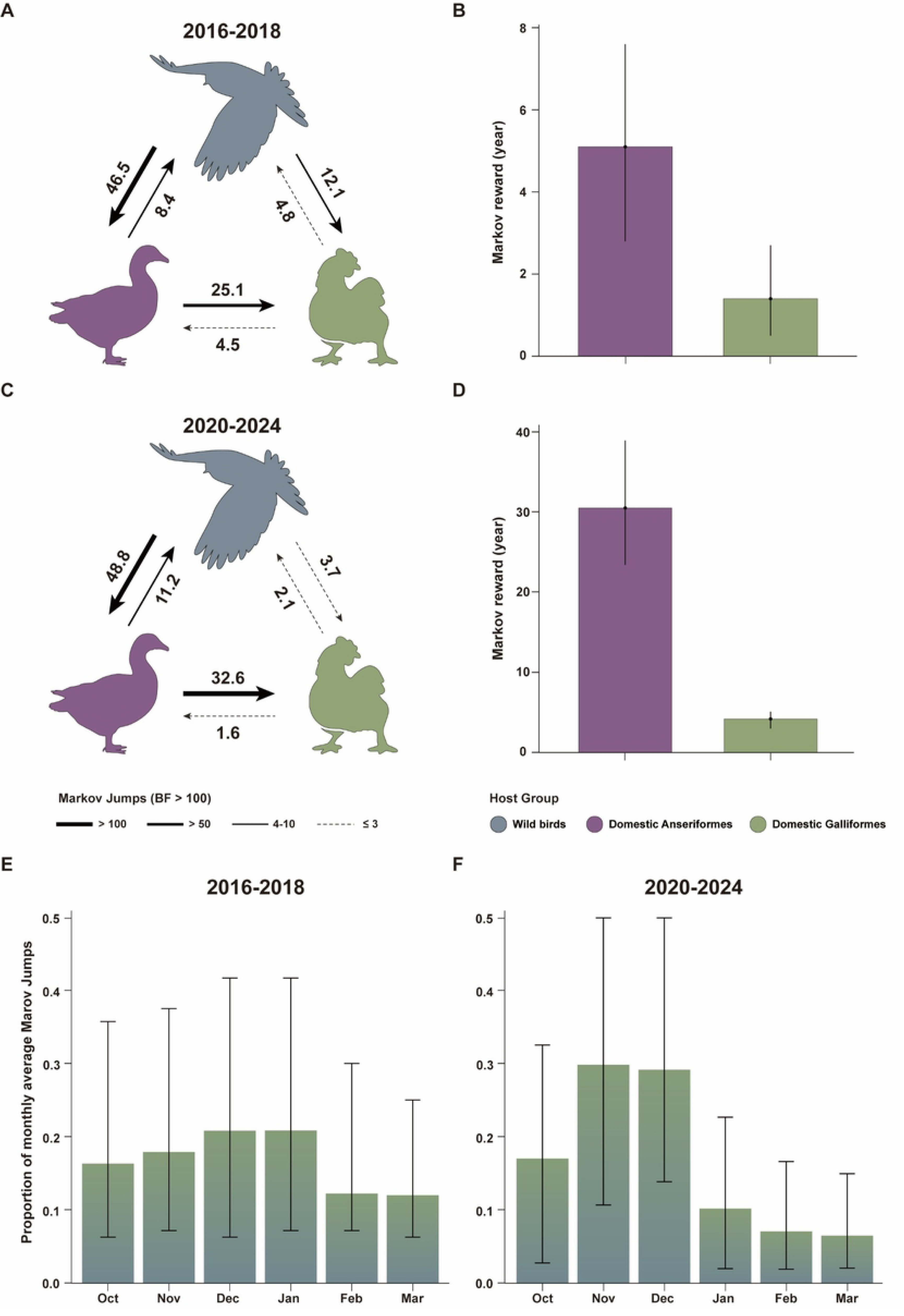
Host-transition dynamics of clade 2.3.4.4b in the ROK across two analysis windows. (A) and (C) Directed host-transition patterns inferred from Markov jump histories. Arrows indicate the direction of inferred host transition, with arrow width proportional to the Bayes factor (BF) for each transition. The numbers next to the arrows represent the fraction of total inferred Markov jumps attributable to each transition. Solid arrows denote supported transitions (BF > 3), whereas dashed arrows denote transitions at BF ≤ 3. (B) and (D) Markov rewards for each host category, shown as posterior mean estimates with 95% HPD intervals. (E) and (F) Monthly distribution of inferred wild-to-poultry transitions (Markov jumps), expressed as the proportion of all inferred transitions in each month. For each posterior draw, monthly transition counts were first converted into proportions by dividing by the total number of wild-to-poultry transitions in that draw. Then posterior medians and 95% credible intervals were calculated across the full set of 1,500 posterior draws as the median and the 2.5th–97.5th percentiles of these draw-level proportions. Results are shown for October 2016–March 2018 in panels (A), (B), and (E), and for October 2020–March 2024 in panels (C), (D), and (F).

### 2.3. Province-level dispersal dynamics

Next, we examined the province-level dispersal of clade 2.3.4.4b using discrete phylogeographic reconstruction (Figure 3 and S6). Transitions from wild birds into JB and JN, and from JN into multiple provinces (including JB, Chungcheongnam-do [CN], and GG), made up a substantial proportion of all inferred province-level transitions (Figure 3). Markov reward analysis suggested that viral lineages were maintained for longer evolutionary periods in JN than in other provinces. Mapping these province-level transition patterns (Figure 4) indicated that wild-to-poultry introductions are disproportionately represented in JN, followed by onward spread to other provinces. Monthly variation in transition shares suggested that inter-province dispersal also exhibited seasonality. Introductions from wild birds into poultry continued into March, albeit at lower frequency. During 2016–2018, onward dispersal from JN frequently followed routes via GG, whereas in 2020–2024 dispersal from JN was more diffuse, with no single dominant route accounting for most transitions.

**Figure 3.**
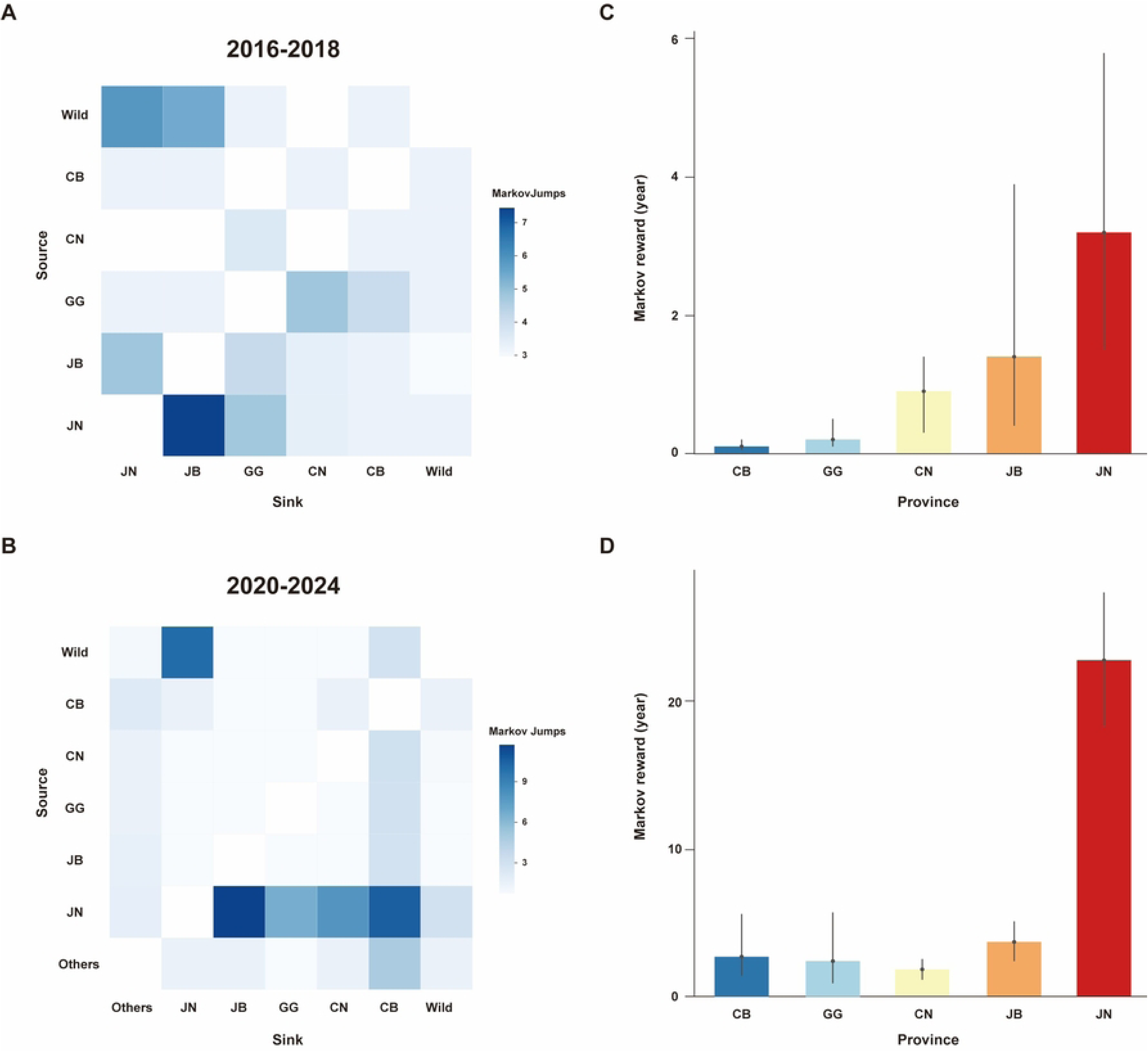
Province-level dispersal dynamics of clade 2.3.4.4b in the ROK across two analysis windows. (A) and (B) Heatmaps summarizing inferred province-to-province transitions from Markov jump histories. Each cell represents a source–sink pair, and values denote the proportion of total inferred Markov jumps attributable to that transition within each analysis window. (C) and (D) Markov rewards for each province, representing inferred lineage residence time, shown as posterior mean estimates with 95% HPD intervals. Results are shown for October 2016–March 2018 in panels (A) and (C), and for October 2020–March 2024 in panels (B) and (D).

**Figure 4.**
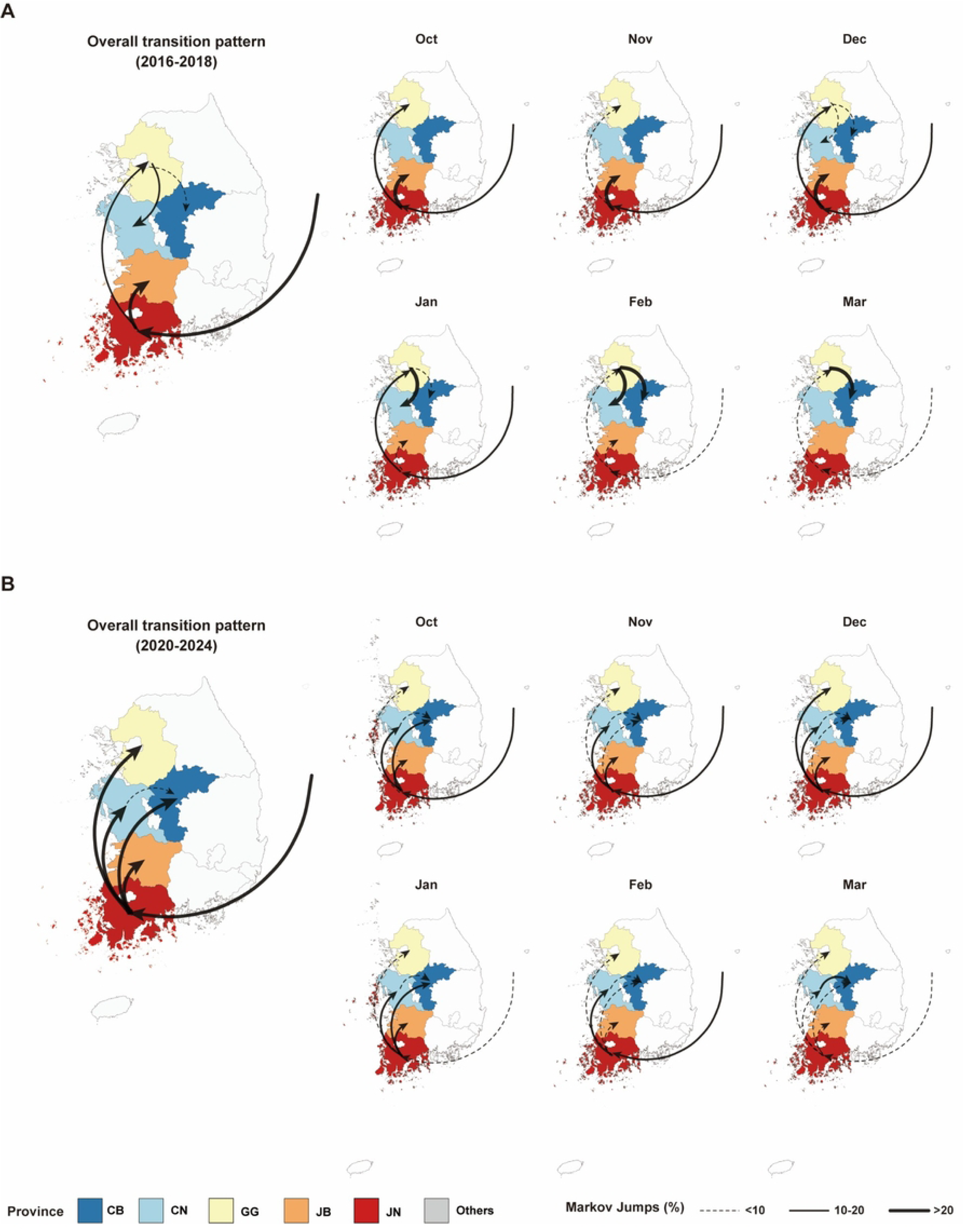
Maps of province-level transition patterns of clade 2.3.4.4b in the ROK across two analysis windows. Maps of inferred inter-province dispersal based on Markov jump histories from the discrete phylogeographic analysis. For each source–sink pair, connections are displayed only when supported by BF > 10, and line width is proportional to the percentage of inferred Markov jumps (%) attributed to that transition. Transitions contributing < 5% (Markov jumps < 5%) are excluded for clarity. The five provinces with the highest outbreak counts are shaded. Each time window is presented as an overall summary (all months combined) and as month-specific summaries (October–March). (A) October 2016–March 2018; (B) October 2020–March 2024. Because sequences of wild birds could not be reliably assigned to specific administrative provinces, the state is represented as an arbitrary reference location in the East Sea to facilitate visualization.

### 2.4. Covariates associated with HPAI dispersal

To identify ecological and epidemiological factors associated with HPAI dispersal, we fitted mixed-effects generalized linear models (GLMs) within the discrete phylogeographic diffusion framework. Summary statistics for each covariate are provided in Table S1. First, we screened covariates using univariable mixed-effects models (Figures S7–S9) and selected the final covariate set to minimize multicollinearity while prioritizing interpretability. Results from the final model are presented in Figure 5. For the wild-to-poultry model, only covariates for the destination province were considered. A significant interaction between the analysis window indicator (2020–2024 vs. 2016–2018) and the number of domestic Anseriformes farms was retained in the final specification. The number of domestic Anseriformes farms showed a significant positive association with the transition rate in both analysis windows, with a modestly stronger effect in 2020–2024. The average probability of AIV infection in wild birds was also positively associated with the transition rate, whereas the 2020–2024 indicator was associated with a small but statistically significant decrease relative to 2016–2018. In this model, province-level random effects were significantly positive for JN and Chungcheongbuk-do (CB), whereas other provinces showed negative or nonsignificant deviations. For the farm-to-farm spread model, covariates for both the origin and destination provinces were considered. The transition rate increased with bigger domestic Galliformes populations and a greater number of domestic Galliformes farms at the destination. Random effects indicated that JN had a significant positive deviation as an origin, while GG showed significant positive deviations as both an origin and a destination. Precision-weighted sensitivity analysis yielded results broadly consistent with the main mixed-effects GLM (Figure S10); however, fixed-effects-only models showed differences in the magnitude and, in some cases, the direction of estimated covariate effects (Figure S11).

**Figure 5.**
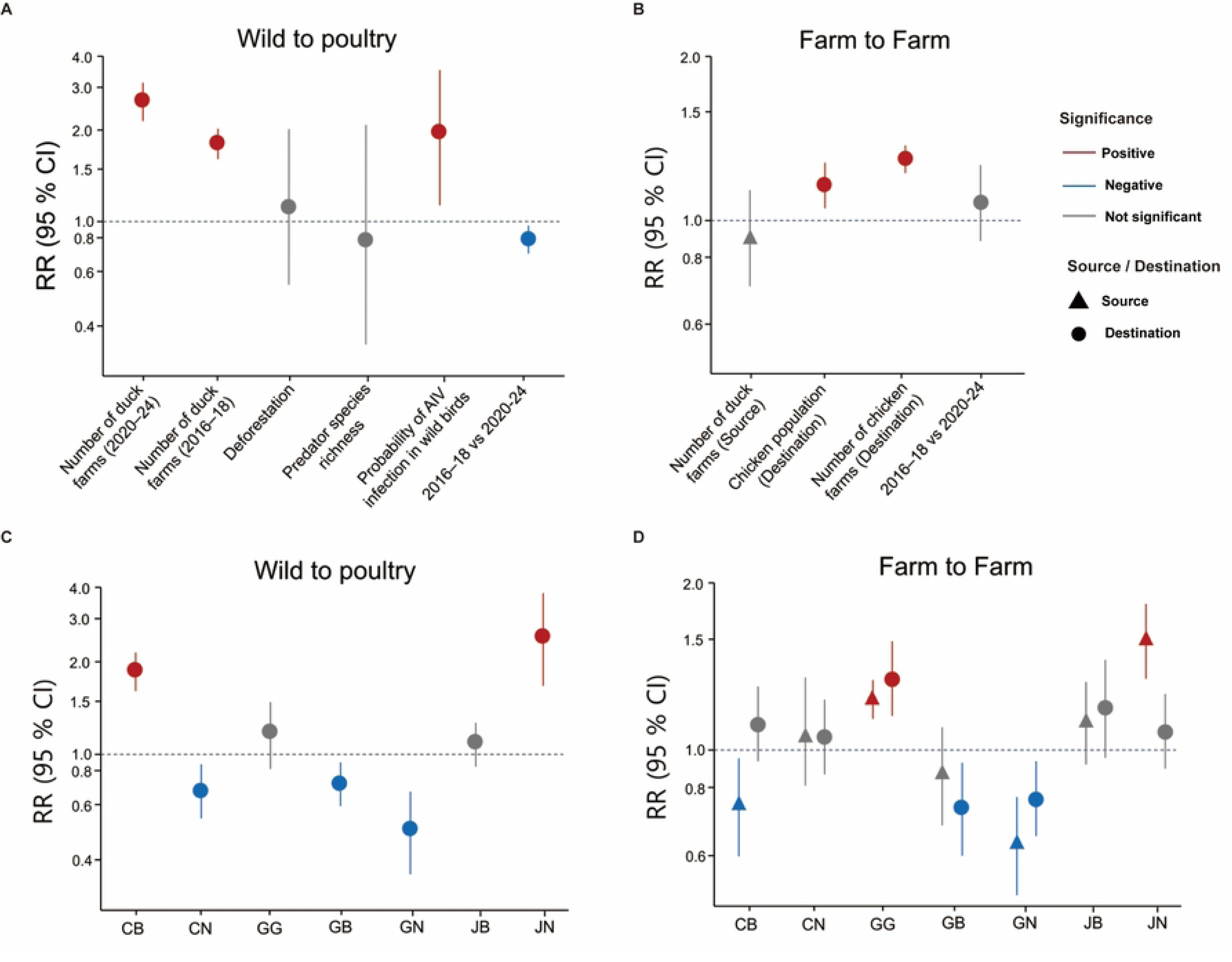
Results of a multivariable mixed-effects GLM for covariates influencing HPAI dispersal. The results are shown for the multivariable mixed-effects models fitted within the phylogeographic diffusion GLM framework. (A) Wild-to-poultry (wild-to-domestic) model: fixed-effect coefficients transformed to rate ratios. (B) Farm-to-farm (domestic-to-domestic) model: fixed-effect coefficients transformed to rate ratios. (C) Wild-to-poultry model: province-specific random effects. (D) Farm-to-farm model: province-specific random effects. For the wild-to-poultry model, only covariates describing the destination province (i.e., where poultry farms are located) were included. For the farm-to-farm model, covariates for both the origin and destination provinces were included.

### 2.5. Sensitivity analyses

Under both downsampling strategies designed to address potential sampling bias (Sensitivity 1: uniform sampling; Sensitivity 2: outbreak-proportional sampling), the inferred phylogeographic patterns were qualitatively consistent with the main analysis (Figures S12–S19). For host-transition dynamics, strongly supported host-transition routes were preserved, and the seasonal timing of wild-to-poultry transitions remained similar. However, Bayesian Markov jump analysis showed that the proportion of transitions attributed to each route shifted somewhat compared to the main analysis. Markov reward analysis indicated a reduced contrast in inferred evolutionary residence times between domestic Anseriformes and Galliformes compared to the main analysis. At the province level, the set of statistically supported dispersal routes and the broad seasonal progression were largely maintained, although relative Markov jump shares and reward allocations showed modest differences. Results from the multivariable mixed effects GLM were also consistent with the main analysis: while effect sizes for some covariates varied, the direction of association and statistical support for key covariates remained unchanged.

## 3. DISCUSSION

Repeated outbreaks of clade 2.3.4.4b viruses in the ROK provide a valuable setting for examining dispersal across the wild–poultry interface and among poultry farms. In this study, we reconstructed their dispersal dynamics, focusing on introductions via wild birds and subsequent transmission among farms. Phylogeographic analyses suggested that domestic Anseriformes farms in the western provinces, particularly Jeollanam-do, likely serve as a key interface for spillover from wild birds into poultry. These introductions tend to occur early in the epidemic season but continue at lower frequency into later months. Transition rates are higher in provinces with more poultry farms, and wild-to-poultry transition rates are greater where the probability of AIV infection in wild birds is elevated. Together, these findings identify the western provinces of the ROK as priority areas for risk-based surveillance and control.

We also found that these waves cluster in spatial hotspots along the western coast. The western provinces of the ROK have consistently been identified as high-risk areas for AIV incursions and subsequent dissemination, not limited to clade 2.3.4.4b (21, 34, 36, 37). This pattern may reflect the presence of suitable habitats for migratory waterbirds (36, 38) as well as the high density of duck farms (37, 39, 40) in these provinces. Migratory waterbirds are widely considered major agents of global AIV dissemination because their high mobility and frequent subclinical infections enable them to carry and shed viruses over long distances (10, 41, 42). The western coast of the ROK features abundant foraging resources for migratory waterbirds, including rice paddies and auxiliary water reservoirs (36, 38, 43). The high density of duck farms in this region may help explain why these areas act as intermediaries between AIV incursions via wild birds and subsequent farm-to-farm spread (34, 37). This pattern may reflect biological characteristics of domestic ducks, which resemble those of wild waterfowl (20, 44) and often show milder clinical signs when infected, potentially facilitating undetected transmission and coinfection with multiple viral strains (45). Collectively, these factors likely increase effective contact at the wild bird–poultry interface, which may serve as an important route of AIV incursion into the ROK.

The recurrent western hotspots also followed a seasonal epidemic cycle. Across five epidemic waves, farm outbreaks typically began in autumn and subsided by the following spring. This pattern suggests that, in the ROK, poultry outbreaks are less consistent with sustained endemic circulation within poultry and instead reflect repeated introductions via wild birds. AIV dispersal by migratory birds is shaped by ecological factors, including migratory flyways and climate (10). For example, Europe has reported seasonal resurgences of infections in wild birds (10), whereas in Taiwan, where clade 2.3.4.4c seems to persist through endemic circulation in poultry, farm outbreaks are observed year-round (13). Although AIV detections in wild birds are reported over a broader temporal window (23), outbreaks on poultry farms exhibit much clearer seasonality. This pattern may reflect resource-driven seasonal shifts in waterfowl movements (36, 46) that increase effective contact between wild birds and poultry, generating peaks in spillover risk. Thus, spillovers into poultry may be governed more by seasonal contact dynamics than by the timing of viral infection in wild birds alone. Contrary to the common expectation that wild-to-poultry transitions are largely confined to the initial phase of an epidemic (32, 34), our analyses indicate that they persist to some extent into the later months of epidemic season. Given that migratory birds enter the ROK via multiple flyways and repeatedly introduce diverse H5 lineages (21, 27, 29), this pattern seems plausible. Collectively, interrupting HPAI epidemics in the ROK will require prioritizing surveillance and control measures in the hotspot regions and sustaining these efforts beyond the early epidemic phase into the later months of the season.

Our phylogeographic GLM analysis provides insight into how ecological and epidemiological factors may shape AIV dispersal in the ROK, where these drivers have received relatively limited attention. A previous study (34) used logistic regression with farm-level HPAI outbreak occurrence as the outcome and reported higher risk under lower temperatures and reduced predator species richness in both chicken and duck farms. It also found elevated risk with greater farm density among chicken farms and with a higher proportion of surrounding nature reserves among duck farms. Although direct comparisons are limited because that study did not incorporate phylogenetic reconstruction and used different covariates, its conclusions are broadly consistent with ours. In our analysis, the number of farms in destination provinces consistently showed a positive association with transition rates, plausibly because more farms increase opportunities for both introduction via wild birds and subsequent farm-to-farm spread. Supporting the importance of this covariate, phylogeographic GLM analyses of AIV dispersal in Taiwan (13) have identified the number of poultry farms as a strongly supported predictor. In addition, the positive association between wild-to-poultry transition rates and the probability of AIV infection in wild birds likely reflects greater introduction pressure from infected wild bird populations. Meanwhile, predator species richness was negatively associated with transition rates in the fixed-effects multivariable model (Figure S11) but became nonsignificant after including province-specific random effects (Figure 5). This attenuation suggests that the apparent protective signal may largely reflect unmeasured province-specific factors captured by the random effects, leaving limited additional explanatory power for predator richness in the main model.

Several locations exhibited positive random-effect deviations (Figure 5), suggesting a residual propensity for transitions not explained by the tested covariates. This pattern likely reflects unmeasured heterogeneity at spatial scales finer than those captured by our province-level covariates, or time-varying processes that are difficult to represent using static predictors. In the wild-to-poultry analysis, the positive deviations observed in CB and JN may indicate elevated incursion propensity at the wild bird–poultry interface, potentially driven by farm-level differences in biosecurity (34) and by fine-scale environmental context, such as proximity to wetlands and variation in local waterfowl activity (36). In the farm-to-farm analysis, JN and GG showed positive random-effect deviations as origins, suggesting that these provinces may act as important sources of onward transmission to other provinces. GG also showed a positive deviation as a destination, indicating that it may play a dual role as both a source and a recipient of viral spread. Consistent with this interpretation, a study of the 2016–2017 H5N6 epidemic in the ROK (32) found that sites linked by poultry production networks or by vehicles involved in farm operations and animal health were more likely to belong to the same phylogenetic clusters, particularly when they shared supply chains. This underscores the role of logistics-mediated contacts in inter-site dissemination. These patterns are compatible with the structure of the Korean poultry industry (47), where vertically integrated production systems are widespread and affiliated farms commonly share supply chains, including feed suppliers, chick sources, and slaughterhouses, thereby intensifying repeated contact. Specifically, JN is a major duck-production hub with clustered facilities, which may amplify onward transmission following incursions from wild birds. GG may be an important conduit to other provinces because of its proximity to the Seoul metropolitan area and its well-developed logistics connectivity, which can facilitate interregional transmission. Together, these findings suggest that unmeasured, province-specific factors may contribute to regional dissemination from and within JN and GG, plausibly driven in part by integrated poultry production networks and associated logistics connectivity.

These findings highlight the value of risk-based control strategies, including targeted quarantine and standstill measures that temporarily halt movements of poultry production vehicles, health-related vehicles, and personnel (32–34). Many of these interventions have been progressively implemented in the ROK following the 2016–2017 AIV epidemic, focusing on high-risk movement pathways and strengthening biosecurity. Consistent with these efforts, our results suggest that farm-to-farm transmission pathways during 2020–2024 were less dominated by a small set of recurrent routes than during 2016–2018 (Figure 4), potentially reflecting disruption of previously persistent dissemination pathways. In the wild-to-poultry analysis, the negative period effect observed for 2020–2024 relative to 2016–2018 (Figure 5) may reflect the implementation of wintertime restrictions on duck production aimed at reducing opportunities for contact between wild birds and poultry (48, 49). Collectively, these patterns suggest measurable benefits from the interventions. However, the association between the number of duck farms and wild-to-poultry transition rates was modestly stronger in 2020–2024 than in 2016–2018, indicating that the concentration of duck farms continues to be a key driver of AIV incursion risk. Consistent with this observation, JN remained a major hotspot for both wild-to-poultry introductions and subsequent farm-to-farm spread. These findings suggest that additional targeted interventions to reduce spillover from wild birds into duck farms may be necessary to further interrupt HPAI epidemics in the ROK.

Sensitivity analyses indicated that statistical support for both inferred transition routes and transition rate–covariate associations was robust to differences in dataset composition. Differences in the distribution of inferred transition events between the main and sensitivity analyses likely reflect shifts in how the model allocates probability when the composition of sampled sequences changes. In the downsampled datasets, the differences in how long lineages were inferred to remain in each category were reduced, likely because sampling was more balanced across categories and therefore suggested smaller gaps in residence times. Some differences between the main analysis and the outbreak-proportional downsampling results may reflect more intensive sampling in provinces with a higher outbreak burden. By contrast, the uniform-sampling results showed qualitative differences, likely reflecting unstable inference due to the very small number of samples included. Overall, the consistency of results across analyses suggests that sampling differences are unlikely to materially affect the main qualitative conclusions.

A key limitation of this study was our reliance on publicly available sequences, which are prone to sampling bias. In the ROK, surveillance of wild birds often depends on voluntary reporting and carcass-based sampling (21), whereas poultry surveillance typically involves proactive sampling at affected sites, leading to systematic differences in sampling intensity across hosts and provinces. To address this issue, we conducted downsampling-based sensitivity analyses using uniform and outbreak-proportional schemes; however, stratified random downsampling may not preserve within-stratum genetic diversity. In principle, subsampling approaches that consider phylogenetic diversity (50) could complement this strategy, but several strata were too sparsely sampled for substantial improvement. To better contextualize potential external sources and introduction pathways into the ROK, we supplemented the dataset with sequences from wild birds from China and Russia. However, this could not fully eliminate residual sampling bias. Improved active surveillance and integration of fine-scale metadata with genomic sequencing is necessary to support more robust inference and risk-based control.

Another limitation was that multiple clade 2.3.4.4b genotypes can be introduced into the ROK within a single season and may differ in transmissibility, host preference, and pathogenicity (21, 23, 27, 29, 30). As a result, joint analyses yield averaged inferences that may obscure genotype-specific dispersal routes and covariate effects. Although HA sequences capture major clade-level dispersal patterns, whole-genome analyses would also be needed to resolve reassortment-driven genotype-specific transmission pathways. Future studies that stratify analyses by genotype or model genotypes as effect modifiers could help clarify genotype-specific drivers and risks.

Finally, the GLM framework that we used had methodological constraints. We applied a two-stage (post hoc) regression approach in which transition rates inferred from Bayesian phylogeographic reconstruction were treated as observed outcomes. This differs from the discrete phylogeographic GLM implemented in BEAST, where transition rates, regression coefficients, and phylogenetic trees are jointly sampled and uncertainty is propagated across model components. Consequently, our approach may not fully propagate uncertainty or capture joint covariance, and the resulting 95% confidence intervals may be narrower than Bayesian highest posterior density intervals. To partially address this limitation, we integrated results across posterior tree samples and assessed robustness using weighted regressions that incorporated uncertainty into transition estimates. Nonetheless, uncertainty may still not be fully captured in the estimated regression coefficients.

Despite these limitations, our study provides a comprehensive reconstruction of the dispersal dynamics underlying recurrent AIV epidemics in the ROK and identifies key factors associated with these patterns. In addition, our findings suggest that risk-based control measures implemented in the ROK may have helped reduce introductions of AIV from wild birds into poultry, while also disrupting long-standing farm-to-farm transmission routes. Accordingly, strengthening risk-based surveillance and control in the western hotspot provinces is essential to mitigate AIV epidemics in the ROK. These efforts should be sustained beyond the early epidemic phase into the later months of the AIV epidemic season.

## 4. MATERIALS AND METHODS

### 4.1. Sequence data preparation

H5 hemagglutinin (HA) sequences of avian influenza A virus clade 2.3.4.4b and associated sample metadata, including collection date, location, and host species, were obtained from the GISAID EpiFlu database (51) on 15 January 2025. We retrieved sequences collected from wild birds and poultry in the ROK between 1 October 2016 and 30 March 2024. The sequences were derived from multiple sampling streams, including cloacal and oropharyngeal swabs, tissues, droppings, and carcass specimens obtained during poultry outbreak investigations and national wild-bird surveillance programs (21, 22). To provide regional context for potential introduction pathways into the ROK and to strengthen phylogenetic reconstruction, we also included sequences from wild birds in China and Russia along major migratory flyways into the ROK (Figure S20) (21, 22, 27). For the non-ROK dataset, stratified random sampling was performed by country and month, selecting up to one sequence per stratum. Duplicate isolates, laboratory-derived isolates, sequences with < 85% HA gene coverage based on the aligned length, sequences containing ≥ 100 ambiguous nucleotides, or those lacking complete collection date information were excluded.

Filtered sequences were aligned using MAFFT v7.490 (52) and trimmed with trimAL v1.4 (53) using a 50% gap threshold, followed by manual trimming to the open reading frame. For quality control prior to time-scaled phylogenetic analyses, maximum-likelihood phylogenies were inferred using IQ-TREE v2.1.4 (54) under the best-fit nucleotide substitution model selected by the ModelFinder algorithm (55). The resulting maximum-likelihood trees were examined for molecular clock outliers using TempEst v1.5.3 (56). Major H5 clades were classified according to the World Health Organization Gs/Gd H5N1 nomenclature (6).

### 4.2. Host and geographic metadata assignment

Host and geographic metadata associated with each sequence were used to assign discrete traits for downstream phylogeographic analyses. Hosts were classified into three groups: wild birds, domestic Anseriformes (primarily domestic ducks), and domestic Galliformes (primarily domestic chickens). Group assignment was based on the GISAID host field and, when necessary, the isolate name. Sequences were retained only when wild/domestic status was explicit or could be inferred from a specified species name; records with ambiguous host annotations (e.g., “avian”) were excluded.

Geographic traits were assigned at the si-do level (the first-level administrative division; hereafter referred to as “provinces”) in the ROK, with the following discrete categories: Wild birds, CB (Chungcheongbuk-do), CN (Chungcheongnam-do), GB (Gyeongsangbuk-do), GG (Gyeonggi-do), GN (Gyeongsangnam-do), JB (Jeollabuk-do), and JN (Jeollanam-do). Other provinces with no eligible poultry sequences were not included as discrete states. Viruses detected in wild birds were treated as a single discrete category regardless of sampling location because some sequences lacked subnational metadata and because key reservoir species are highly mobile, that is, the sampling location may not reflect the location where transmission is sustained (19, 31, 57). For domestic poultry sequences with only country-level location metadata, province assignments were resolved by matching collection date and poultry type with publicly available official Korean HPAI outbreak reports from the Korea Animal Health Integrated System (http://www.kahis.go.kr). These surveillance records include farm location at the eup-myeon-dong level (the third-level administrative division), outbreak date, poultry type, outbreak size, and the diagnosing laboratory. We first searched for reports within ±3 days of the collection date with the same poultry type; if none were found, the window was expanded to ±7 days. Then the province from the matched report was assigned to the sequence. If multiple reports matched, the province was assigned only when all matched reports corresponded to the same province; otherwise, the sequence was excluded.

Following these procedures, the primary analytical dataset included 419 sequences: 121 from wild birds from the ROK, 62 from wild birds from China and Russia, and 236 from domestic poultry from the ROK.

### 4.3. Bayesian phylogenetic reconstruction

The study period was divided into two distinct windows (October 2016–March 2018 and October 2020–March 2024) to align with the typical autumn–spring migratory season (approximately October–March) (36), when poultry outbreaks are most frequent. Although poultry farm outbreaks associated with the 2.3.4.4b clade are generally considered to have begun in autumn 2017, introductions of this clade into wild birds were inferred to predate this period. Accordingly, we set the start of the study period to October 2016 to account for this earlier circulation. This design was also motivated by the near absence of clade 2.3.4.4b introductions and the lack of reported poultry farm outbreaks during the intervening period (April 2018–September 2020). Thus, we excluded this interval to avoid conflating temporally disconnected epidemic phases. Bayesian phylogenetic reconstruction and all downstream analyses were performed separately for each window.

Bayesian phylogenetic reconstructions based on Markov chain Monte Carlo (MCMC) sampling were conducted in BEAST v1.10.4 (58) with computational acceleration via the BEAGLE library (59). The best-fitting nucleotide substitution model was identified using ModelFinder as implemented in IQ-TREE v2.1.4, which selected GTR+I+G4. Accordingly, we used a general time-reversible (GTR) substitution model with an estimated proportion of invariant sites (+I) and among-site rate heterogeneity modeled using a discrete gamma distribution with four categories (+G4). Rate heterogeneity across branches was modeled using an uncorrelated lognormal relaxed molecular clock (60), and population history was modeled using a constant-size coalescent tree prior (61) with a lognormal prior on the effective population size.

For each window-specific dataset, we ran at least four independent MCMC chains for 100 million states, sampling every 10,000 states and removing the first 10% as burn-in. Convergence and mixing were assessed in Tracer v1.7.1 after combining log files using LogCombiner, and all parameters were required to achieve effective sample size values > 200. When necessary, we increased the burn-in fraction and/or ran additional chains until the effective sample size criterion was met. The posterior tree files from the retained chains were combined, and trees were thinned at regular intervals to obtain an empirical posterior distribution of 1,500 trees per analysis window to reduce computational burden for downstream phylogeographic analyses (10). The MCC tree was subsequently summarized from this empirical set using TreeAnnotator.

### 4.4. Discrete trait diffusion models

To investigate viral spread among administrative regions and avian host groups, we implemented a discrete-trait diffusion model in BEAST v1.10.4. This framework models transitions among discrete states along the phylogeny as a continuous-time Markov chain and enables reconstruction of ancestral trait histories (62). In the discrete-trait analysis, we used an asymmetric substitution model to estimate directional diffusion rates between provinces. Transition rates were estimated under a Bayesian stochastic search variable selection (BSSVS) procedure. BFs were computed from the BSSVS output using SpreaD3 v0.9.6 (63) to quantify statistical support for individual diffusion routes. Diffusion routes were considered statistically supported at BF > 3.0, strongly supported at BF > 10, and decisively supported at BF > 100 (42). Markov jump transitions between states were summarized by their posterior expected counts and, where relevant, normalized as proportions of all inferred transitions across the phylogeny. Markov rewards were estimated to quantify the proportion of time spent in each discrete state (64).

We translated Markov jump times to calendar dates using the matched time-scaled tree associated with each complete-history log. Jump times were recorded as years before present and converted into calendar dates by anchoring time 0 to the most recent collection date within each analysis window (62, 64). Mapped events were subsequently summarized at monthly and yearly resolution. All post hoc processing of mapped Markov jumps, together with result visualization and map generation, was performed in R v4.2.1 (R Core Team).

### 4.5. Discrete phylogeographic Generalized Linear Model (GLM) analysis

To evaluate covariates associated with discrete-trait transition rates among provinces, we performed a post hoc mixed-effects regression on posterior samples of transition rates inferred under a discrete phylogeographic diffusion model. Conceptually, this approach parallels the discrete phylogeographic GLM implemented in BEAST (65) and post hoc landscape phylogeographic analyses in continuous phylogeography (66, 67). Accordingly, we fitted a log-linear regression model for transition rates and propagated uncertainty using posterior samples. Because the standard discrete phylogeographic GLM in BEAST imposes a single coefficient vector across ordered transition pairs, we adopted a post hoc framework in which covariate effects were allowed to vary by transition type, thereby accommodating mechanistic differences between wild-to-poultry spillover and farm-to-farm spread.

Specifically, we extracted instantaneous transition rates (*q*) for each MCMC draw from an empirical posterior sample of trees. We considered 24 candidate covariates (Table S2), all of which were log-transformed and standardized. To allow mechanistically distinct covariate effects, transition rates were analyzed separately by transition type, thereby permitting type-specific coefficient vectors: wild-to-poultry spillover (wild birds → provinces; unidirectional) and farm-to-farm spread (province → province; bidirectional). For the first one, covariates were defined for destination provinces only. To propagate uncertainty in the regression, draw-specific rates were treated as repeated measurements, with a draw-level random intercept included to capture within-draw dependence. Posterior rate samples were pooled across the two analysis windows, and a fixed effect for the window was included to account for between-window differences. As sensitivity analyses, we compared the mixed-effects GLM results with alternative model specifications, including precision-weighted and fixed-effects-only models. Full model specifications and implementation details are provided in the Supplementary Methods.

### 4.6. Sensitivity analyses for sampling bias

To evaluate the robustness of phylogeographic inference to sampling bias, we conducted two downsampling-based analyses after stratifying sequences by time window and discrete-trait state. These sensitivity analyses were performed separately for the geographic diffusion analysis and for the host-group diffusion analysis. In Sensitivity 1, each stratum was randomly downsampled to an equal number of sequences, with the target per stratum set to the minimum number of sequences observed across strata within the same time window. In Sensitivity 2, stratum-specific targets were made proportional to reported poultry outbreak counts by time window, province, and poultry type. Because comparable incidence data were unavailable for the wild-bird state, we assigned the wild-bird stratum a target equal to the mean target across domestic poultry strata within the same time window.

In both sensitivity analyses, when one or more strata could not reach the target because of limited sequence availability, a common downscaling (capping) rule was applied so that final targets did not exceed the number of sequences available in any stratum. For each sensitivity dataset, we reran the full time-scaled Bayesian phylogenetic reconstruction and discrete-trait diffusion inference in BEAST and repeated the downstream analyses using the same workflow as in the main analysis. The resulting sample compositions for these analyses are summarized in Tables S3 and S4.

## Conflicts of Interest

The authors declare that there are no conflicts of interest regarding the publication of this article.

## Ethics statement

This study used publicly available viral sequence data and publicly reported animal outbreak and surveillance metadata. No new animal sampling, animal experiments, or human participant data collection was conducted as part of this study; therefore, ethical approval was not required.

## Authors’ contributions

TC conceived the study, analyzed the data, and wrote the manuscript. SL contributed to data analysis and interpretation. JIK contributed to data interpretation and was involved in the manuscript’s review and revision process. K-DM was responsible for the conception of the study, assisted in data interpretation, and was involved in the manuscript’s review and revision process. All authors provided input on the study’s design and data analysis and made significant contributions to the discussion and interpretation of the results.

## Acknowledgements

This research was supported by a National Research Foundation of Korea (NRF) grant funded by the Korea government (MSIT) (grant number: RS-2025-00517731). However, the funders had no role in study design or conduct, data collection, management, analysis, or interpretation, manuscript preparation, review, or approval, or the decision to submit the article for publication.

## Availability of data and materials

The data sources used in this study are described in the Materials and Methods and Supplementary Materials. Raw sequence data were obtained from the GISAID EpiFlu database and are available from GISAID subject to its data access terms. Publicly available outbreak and surveillance data sources used in the analysis are described in the Materials and Methods and Supplementary Materials. The analysis code necessary to reproduce the analyses is available at the following repository: https://github.com/TaeHChang/Spatiotemporal-Dynamics-of-Highly-Pathogenic-Avian-Influenza-H5-Virus-.

## Notes

### Competing Interest Statement

The authors have declared no competing interest.

